# SARS-CoV-2-induced humoral immunity through B cell epitope analysis and neutralizing activity in COVID-19 infected individuals in Japan

**DOI:** 10.1101/2020.07.22.212761

**Authors:** Shota Yoshida, Chikako Ono, Hiroki Hayashi, Satoshi Shiraishi, Kazunori Tomono, Hisashi Arase, Yoshiharu Matsuura, Hironori Nakagami

## Abstract

The aim of this study is to understand adaptive immunity to SARS-CoV-2 through the analysis of B cell epitope and neutralizing activity in coronavirus disease 2019 (COVID-19) patients. We obtained serum from thirteen COVID-19 patients. Most individuals revealed neutralizing activity against SARS-CoV-2 assessed by a pseudotype virus-neutralizing assay. The antibody production against the spike glycoprotein (S protein) or receptor-binding domain (RBD) of SARS-CoV-2 was elevated, with large individual differences, as assessed by ELISA. In the analysis of the predicted the linear B cell epitopes, two regions (671-690 aa. and 1146-1164 aa.), which were located in S1 and S2 but not in the RBD, were highly reactive with the sera from patients. In the further analysis of the B cell epitope within the S protein by utilizing a B cell epitope array, a hot spot in the N-terminal domain of the S protein but not the RBD was observed in individuals with neutralizing activity. Overall, the analysis of antibody production and B cell epitopes of the S protein from patient serum may provide a novel target for the vaccine development against SARS-CoV-2.

## Introduction

The recent emergence of severe acute respiratory syndrome coronavirus 2 (SARS-CoV-2) and the resulting coronavirus disease 2019 (COVID-19) poses an unprecedented health crisis that was declared a pandemic by the World Health Organization (WHO) (Wang *et al*, 2020). To fight against COVID-19, the rapid development of a vaccine is required. SARS-CoV-2 belongs to the Betacoronavirus genus, and SARS-CoV-1 and Middle East respiratory syndrome coronavirus (MERS-CoV) are two highly pathogenic viruses (Lu *et al*, 2020; Chan *et al*, 2020; Wu *et al*, 2020a). The spike glycoprotein (S) on the SARS-CoV-2 surface plays an essential role in receptor binding and virus entry, and previous studies on SARS-CoV-1 and MERS-CoV have revealed the importance of the S protein as a potential antigen target for vaccines (Lu *et al*, 2015; Wu *et al*, 2020b; Hoffmann *et al*, 2020). The S protein has been found to induce robust and protective humoral and cellular immunity, including the development of neutralizing antibodies and T cell-mediated immunity (Vabret *et al*, 2020; Du *et al*, 2009).

To understand the immune response to COVID-19, the analysis of virus-specific CD4^+^ and CD8^+^ T cells is required. Grifoni *et al* (2020b) recently demonstrated that using HLA class I and II predicted peptide ‘megapools’, circulating SARS-CoV-2-specific CD8^+^ and CD4^+^ T cells were identified in ~70% and 100% of COVID-19 convalescent patients, respectively. CD4^+^ T cell responses to S protein were robust and correlated with the magnitude of the anti-SARS-CoV-2 IgG and IgA titers. Importantly, the antibody titer for the receptor-binding domain (RBD) of the S protein correlated well with an increase in spike-specific CD4^+^ T cell responses but not non-Spike-specific CD4^+^ T cell responses. In other reports, RBD-specific antiviral T cell responses have also been detected in people who have recovered from COVID-19 (Vabret *et al*, 2020).

Here, we addressed the humoral immune response by measuring antibody production against S protein and the neutralizing ability in convalescent patients from two different hospitals. In addition, the B cell epitope of S protein was analyzed by peptide epitope array. These results will assist vaccine design and evaluation of candidate vaccines.

## Results

### Antibody production and neutralizing activity in thirteen serum samples

To investigate the humoral immunoreaction to SARS-CoV-2, we assessed 13 serum samples collected from COVID-19 patients. Out of 13 patients, seven patients were in the intensive care unit of Osaka University Hospital (OU samples), and six patients were in Osaka City Juso Hospital (Ju samples). To estimate the existence of antibodies against SARS-CoV-2, we performed 75% plaque-reduction neutralization tests (PRNT75) using pseudotyped VSVs. At an evaluation point of the PRNT75 (Fig 1 and Table 1), we confirmed neutralizing activity in most of the samples, especially high in OU #2, #3, and #6; however, in a few samples (OU #5 and #7 and Ju #3), neutralizing activity was not detected. We speculate that the disease phase and severity of patients may be correlated with these neutralizing activities because most of the patients in Osaka University Hospital are treated in the intensive care unit (ICU) and are more severe than those in Juso Osaka City Hospital.

**Figure 1.**
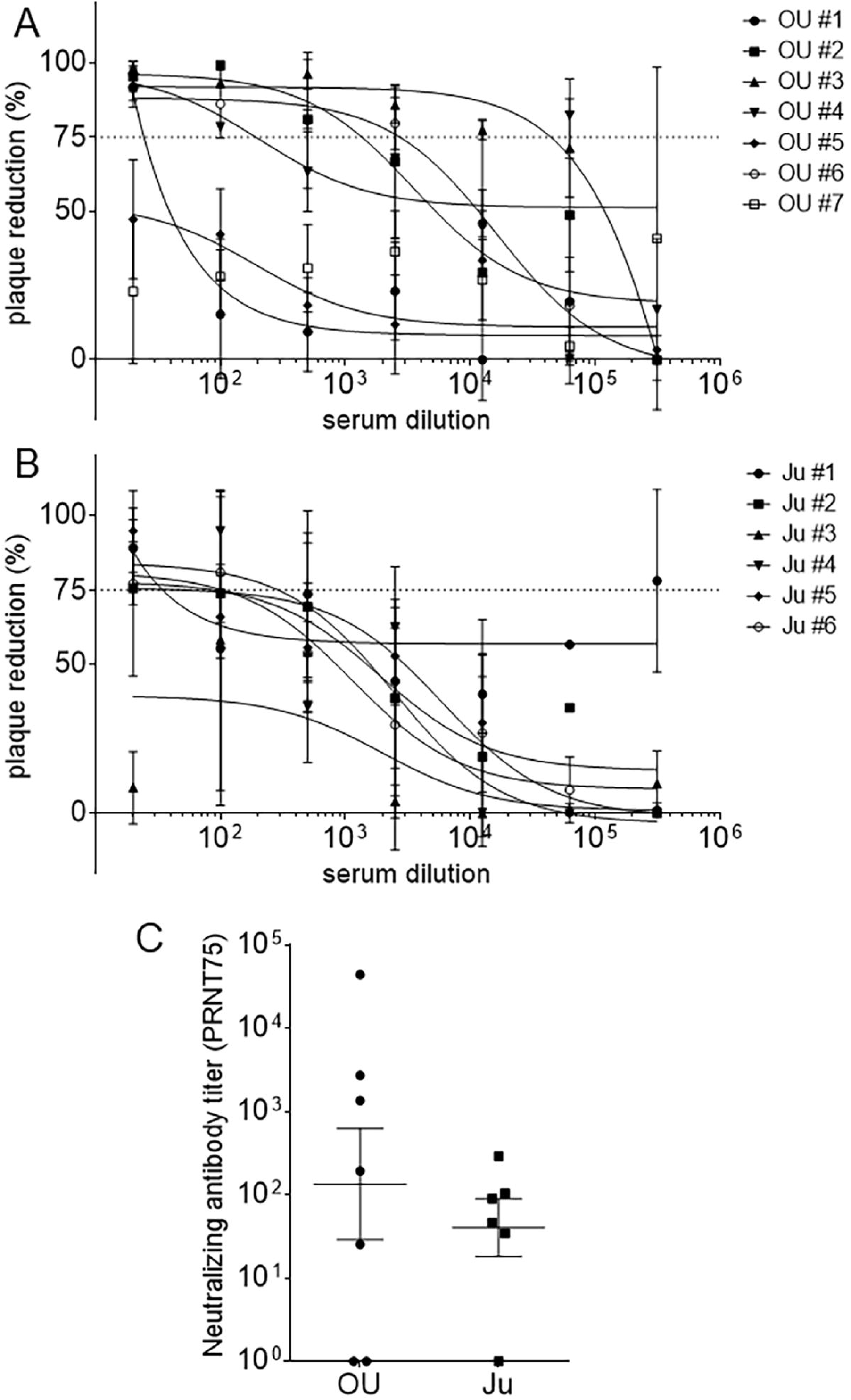
Plaque-reduction neutralization test for serum samples of COVID-19 patients. **(A and B)** The sigmoidal dose-response curve of neutralizing activities of serum antibodies against SARS-CoV-2 with a PRNT75. **(A)** OU: serum samples collected from patients in the ICU of Osaka University Hospital. **(B)** Ju: serum samples collected from patients in Osaka City Juso Hospital. (C) Neutralizing antibody titers of serum samples at an evaluation point of the PRNT75. All the data are expressed as the mean ± SEM. Statistical evaluation was performed by unpaired two-tailed t-test (**C**, natural logarithmic transformation).

**Table 1.** Comparison of several antibody test kits as a screening test for the SARS-CoV-2-specific IgG or IgM.

| Sample | KURABO |  | Vazyme |  | Abnova |  | Raybiotech |  | PRNT<br>75% titer |
| --- | --- | --- | --- | --- | --- | --- | --- | --- | --- |
|  | IgM | IgG | IgM | IgG | IgM | IgG | IgM | IgG |  |
| OU #1 | — | + | — | + | — | + | — | — | 24.4 |
| OU #2 | + | + | + | + | + | + | + | + | 1356.2 |
| OU #3 | — | + | — | + | + | + | + | + | 44126.6 |
| OU #4 | — | + | — | + | + | + | + | + | 192.0 |
| OU #5 | — | — | — | — | — | — | — | — | — |
| OU #6 | — | + | — | + | + | + | + | + | 2727.8 |
| OU #7 | + | + | — | + | + | + | + | — | — |
| Ju #1 | — | + | — | + | + | + | + | + | 34.4 |
| Ju #2 | — | + | — | + | + | + | + | + | 89.0 |
| Ju #3 | + | — | — | + | — | + | + | + | — |
| Ju #4 | + | + | + | + | — | + | + | + | 289.4 |
| Ju #5 | + | + | — | + | — | + | + | — | 45.4 |
| Ju #6 | + | + | — | + | + | + | + | + | 103.3 |
| Negative | — | — | — | — | — | — | — | — | — |

OU, serum samples collected from patients in the ICU of Osaka University Hospital.
Ju, serum samples collected from patients in Osaka City Juso Hospital.
Negative, negative control serum.
PRNT, Plaque-reduction neutralization test.

We also tested serum samples with several antibody test kits as a screening test for SARS-CoV-2-specific IgG or IgM antibodies. Table 1 shows that most of the samples tested positive for IgG and were correlated with neutralizing antibody activity, except for a few samples (OU #7 and Ju #3). The results of the IgM antibody test, in contrast, varied among antibody test kits.

In the analysis of the humoral response to SARS-CoV-2, we focused on portions of S protein, such as the S1 subunit, S2 subunit, and RBD in the S1 subunit, as candidate antigens. ELISA showed that several sera collected from COVID-19 patients strongly reacted with SARS-CoV-2 recombinant proteins (Table 2 and Appendix Table S1), and we selected spike S1+S2 recombinant protein from Beta Lifescience and RBD recombinant protein from Beta Lifescience for further experiments. As shown in Fig 2A and 2B, increased antibody titers to spike (S1+S2) IgG were observed in most of the COVID-19 patients; however, this increase was not observed in a few patients. Interestingly, one sample (#6) from a patient in Osaka University Hospital indicated an extremely high titer of antibody. The average antibody titer of the patients in Osaka University Hospital was higher than that of the patients in Juso Osaka City Hospital. We speculate that the disease phase and severity of patients may be also correlated with the total anti-spike IgG titer because most of the patients in Osaka University Hospital were treated in the ICU and were more severe than those in Juso Osaka City Hospital. We additionally analyzed the IgM and IgG subclasses of anti-spike (S1+S2) antibodies. IgG1 was mainly detected, and IgG3 was less detected (Fig 2C and 2D). Neither IgG2 nor IgG4 were detected (Appendix Fig S1A and S1B). Compared with the IgG titer, the IgM titer was much lower in all of the patients (Fig 2E and 2F) because all of the samples were obtained from the patients in the recovery phase, not in the acute phase.

**Table 2.** Screening of recombinant proteins for ELISA to measure anti-S1, S2, S1+S2 or RBD antibodies.

| Sample | Sino Biological |  | Beta Lifescience |  |  |  | Ray Biotech |  |
| --- | --- | --- | --- | --- | --- | --- | --- | --- |
|  | S1 | S2 | S1 | S2 | S1+S2 | RBD | S1 | S2 |
| OU #1 | 3.5* | 3.5* | 3.5* | 2.548 | 2.664 | 2.238 | 2.742 | 3.5* |
| OU #2 | 3.5* | 3.5* | 3.5* | 2.644 | 2.792 | 3.286 | 3.004 | 3.5* |
| OU #3 | 3.5* | 3.5* | 3.5* | 2.461 | 2.549 | 2.744 | 1.263 | 3.5* |
| OU #4 | 0.624 | 3.5* | 0.366 | 1.262 | 0.519 | 0.401 | 0.186 | 1.883 |
| OU #5 | 0.07 | 3.5* | 0.076 | 0.236 | 0.177 | 0.102 | 0.153 | 0.427 |
| Negative | 0.154 | 3.5* | 0.16 | 0.124 | 0.129 | 0.121 | 0.265 | 0.188 |

OD at 450 nm, 10-fold dilution.
3.5\*, over 3.5 Optical Density (calculated as 3.5)

**Figure 2.**
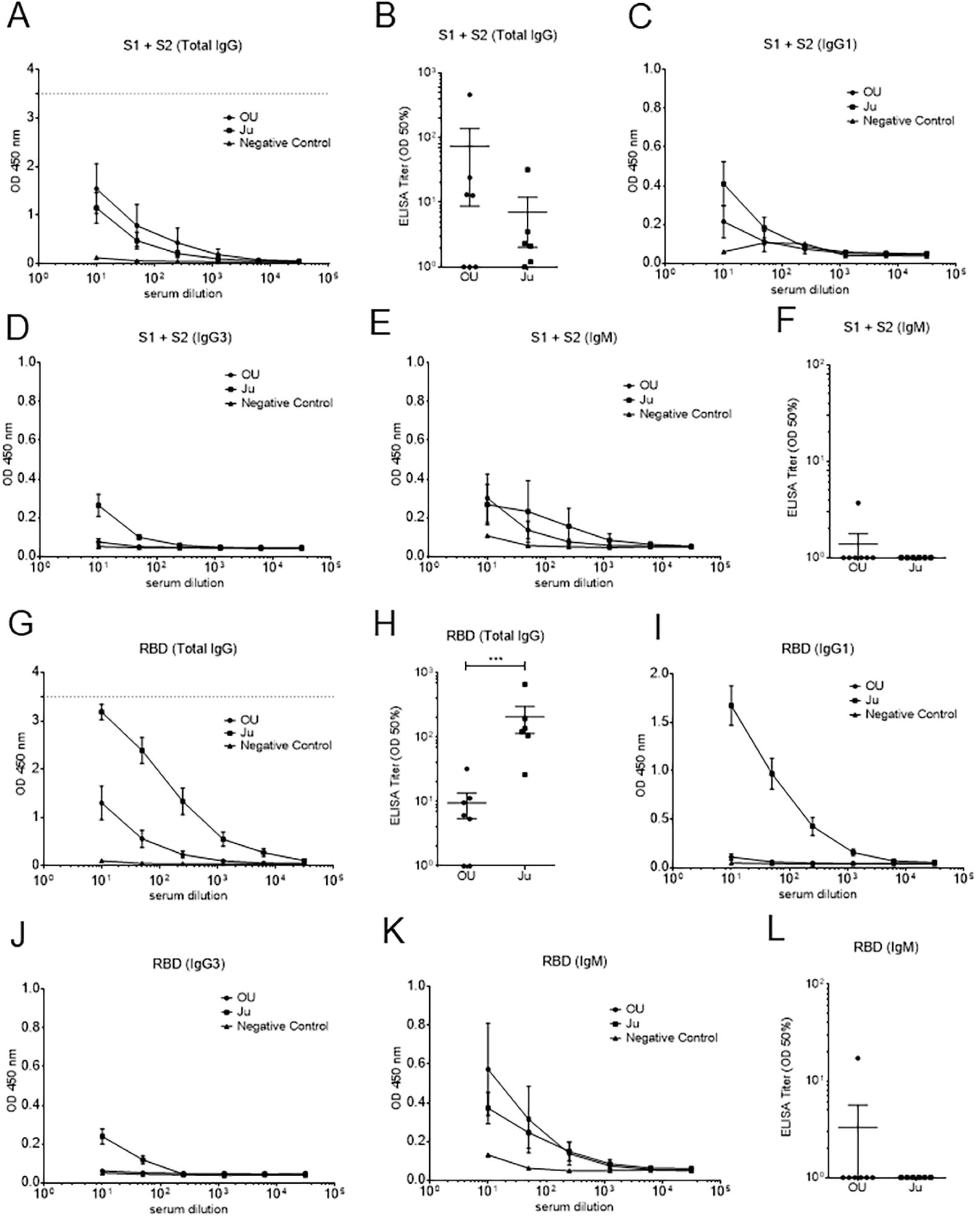
Anti-SARS-CoV-2 IgG, IgM, and IgG subclass responses of COVID-19 patients. **(A-F)** The serum titer against recombinant SARS-CoV-2 spike S1+S2 protein. **(A)** Total IgG, **(C)** IgG1, **(D)** IgG3, and **(E)** IgM, expressed as the OD at 450 nm. **(B)** Total IgG and **(F)** IgM, expressed as the half-maximal binding (OD 50%). **(G-L)** The serum titer against recombinant SARS-CoV-2 spike RBD protein. **(G)** Total IgG, **(I)** IgG1, **(J)** IgG3, and **(K)** IgM, expressed as the OD at 450 nm. **(H)** Total IgG and **(L)** IgM, expressed as the OD 50%. OU, serum samples collected from patients in the ICU of Osaka University Hospital; Ju, serum samples collected from patients in Osaka City Juso Hospital. All the data are expressed as the mean ± SEM. Statistical evaluation was performed by unpaired two-tailed t-test (**B, F, H and L**, natural logarithmic transformation); ***p < 0.001.

Since the RBD in the S1 subunit is the major target for neutralizing antibodies, we next focused on anti-spike RBD antibodies. Similarly, an increased IgG titer but not IgM titer was observed in most of the samples (Fig 2G, 2H, 2K, and 2L). Interestingly, the average anti-spike RBD IgG titer of the patients in Osaka University Hospital was lower than that of the patients in Juso Osaka City Hospital, which was the opposite result from the anti-spike (S1+S2) IgG titer. We speculate that anti-RBD antibody titers may be correlated with good progression because some of the patients in Juso Osaka City Hospital recovered and were in the convalescent phase. In terms of IgG subclasses, IgG1 was mainly detected, and IgG3 was less detected (Fig 2I and 2J), but neither IgG2 nor IgG4 were detected (Appendix Fig S1C and S1D).

### Analysis of B cell epitopes in S protein in thirteen serum samples

Furthermore, we predicted linear B cell epitopes by BepiPred-2.0, a prediction tool of B cell epitopes assessed by the characterization of amino acid sequences. Five regions (AH-531, 346-365 aa.; AH-526, 413-432 aa.; AH-527, 442-460 aa.; AH-532, 491-509 aa.; and AH-533, 518-537 aa.) were selected from the RBD, and three regions (AH-528, 146-164 aa.; AH-529, 671-690 aa.; and AH-530, 1146-1164 aa.) were selected from other regions (Fig 3A and Appendix Table S2). In the analysis of IgG (Fig 3B), two regions, AH-529 (including the S1/S2 cleavage site) and AH-530 (heptad repeat 2, HR2), reacted with sera from most of the patients. Four regions of the RBD (AH-526, AH-527, AH-532 and AH-533) mildly reacted with sera from some of the patients, and the other regions, AH-528 (N-terminal domain, NTD) and AH-531 (RBD), did not react. In the analysis of IgM (Fig 3C), in contrast, four regions (AH-526, AH-527, AH-529, and AH-530) mildly reacted with sera from some of the patients, and the other four regions hardly reacted.

**Figure 3.**
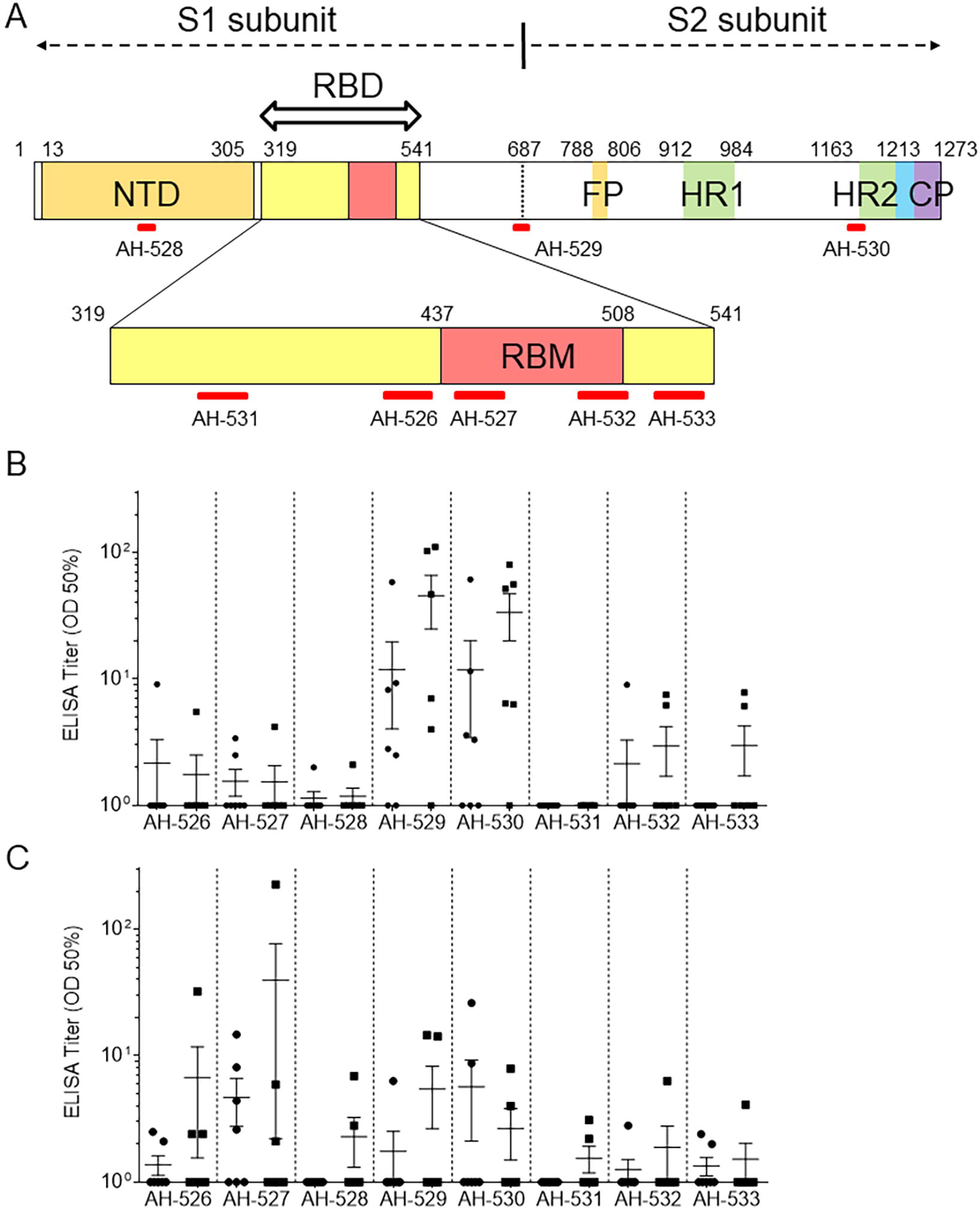
Linear B cell epitopes on Spike protein of SARS-CoV-2 determined by ELISA. **(A)** Scheme of predicted linear B cell epitopes in the RBD or other regions of Spike protein in SARS-CoV-2. As shown in red bars, five regions (AH-531, 346-365 aa.; AH-526, 413-432 aa.; AH-527, 442-460 aa.; AH-532, 491-509 aa.; and AH-533, 518-537 aa.) were selected from the RBD, and three regions (AH-528, 146-164 aa.; AH-529, 671-690 aa.; and AH-530, 1146-1164 aa.) were selected from the NTD, S1 and S2 subunits. NTD, N-terminal domain; RBD: receptor-binding domain; RBM, receptor-binding motif; FP, fusion peptide; HR1, heptad repeat 1; HR2, heptad repeat 2; CP, cytoplasm domain. **(B and C)** The serum titer against synthetic SARS-CoV-2 peptides (AH-526 to AH-533) is expressed as the half-maximal binding (OD 50%). **(B)** Total IgG and **(C)** IgM. closed circle, serum samples collected from patients in the ICU of Osaka University Hospital; closed square, serum samples collected from patients in Osaka City Juso Hospital. All the data are expressed as the mean ± SEM.

We further evaluated the linear B cell epitope within spike protein using a CelluSpots peptide array composed of a series of 15-mer peptides overlapping by five amino acids (i.e., 1-15 aa., 5-20 aa., 10-25 aa., etc.). The lists and maps of the top 20 peptides with high intensity values for each sample (Fig 4 and Table 3) show that a large number of strongly binding B cell epitopes were located in the regions outside the RBD, such as the NTD, fusion peptide (FP), HR2 and cytoplasm domain (CP). For instance, several strongly binding epitopes in samples OU #1, #2, and #6 were located in the CP, NTD, and FP, respectively. Moreover, we evaluated the linear B cell epitope within nucleocapsid, membrane and envelope proteins using a CelluSpots peptide array. As shown in Appendix Table S3, most of the strongly binding B cell epitopes were located in nucleocapsid protein.

**Figure 4.**
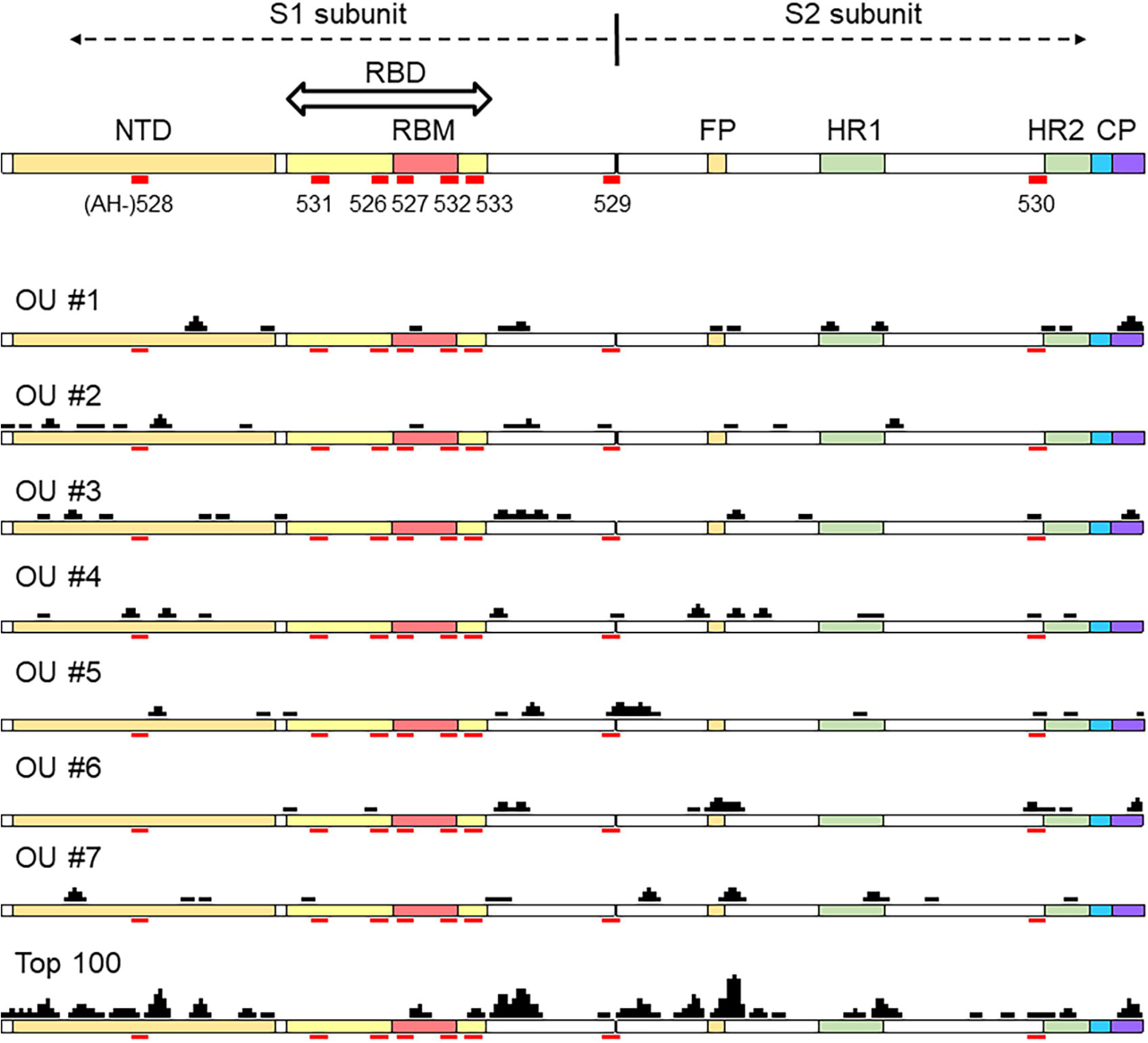
Mapping of the top 20 strongest binding peptide regions of serum samples. The scheme of lists and maps of the top 20 peptide regions with high intensity values for each individual (OU #1-#7). A series of 15-mer peptides overlapping by five amino acids (i.e., 1-15 aa., 5-20 aa., 10-25 aa., etc.) were displayed in this peptide array. The black block indicates the top 20 strongest binding peptide regions with serum samples from each individual and the top 100 strongest binding peptide regions for all samples. The red bars show the candidate linear B cell epitopes: five regions from the RBD (AH-531, 346-365 aa.; AH-526, 413-432 aa.; AH-527, 442-460 aa.; AH-532, 491-509 aa.; and AH-533, 518-537 aa.) and three regions from the NTD, S1 and S2 subunits (AH-528, 146-164 aa.; AH-529, 671-690 aa.; and AH-530, 1146-1164 aa.). NTD, N-terminal domain; RBD: receptor-binding domain; RBM, receptor-binding motif; FP, fusion peptide; HR1, heptad repeat 1; HR2, heptad repeat 2; CP, cytoplasm domain.

**Table 3.** Top 20 strongest binding peptide regions of serum samples from each individual (OU #1-7)

| #1 | Epitope ID | Position | Amino Acid Sequence |
| --- | --- | --- | --- |
| 1 | K12 | 1256 - 1270 | F-D-E-D-D-S-E-P-V-L-K-G-V-K-L |
| 2 | K11 | 1251 - 1265 | G-S-C-C-K-F-D-E-D-D-S-E-P-V-L |
| 3 | I 3 | 971 - 985 | G-A-I-S-S-V-L-N-D-I-L-S-R-L-D |
| 4 | J17 | 1161 - 1175 | S-P-D-V-D-L-G-D-I-S-G-I-N-A-S |
| 5 | B20 | 216 - 230 | L-P-Q-G-F-S-A-L-E-P-L-V-D-L-P |
| 6 | K10 | 1246 - 1260 | G-C-C-S-C-G-S-C-C-K-F-D-E-D-D |
| 7 | E19 | 571 - 585 | D-T-T-D-A-V-R-D-P-Q-T-L-E-I-L |
| 8 | D20 | 456 - 470 | F-R-K-S-N-L-K-P-F-E-R-D-I-S-T |
| 9 | E20 | 576 - 590 | V-R-D-P-Q-T-L-E-I-L-D-I-T-P-C |
| 10 | G19 | 811 - 825 | K-P-S-K-R-S-F-I-E-D-L-L-F-N-K |
| 11 | I 4 | 976 - 990 | V-L-N-D-I-L-S-R-L-D-K-V-E-A-E |
| 12 | C11 | 291 - 305 | C-A-L-D-P-L-S-E-T-K-C-T-L-K-S |
| 13 | B19 | 211 - 225 | N-L-V-R-D-L-P-Q-G-F-S-A-L-E-P |
| 14 | E16 | 556 - 570 | N-K-K-F-L-P-F-Q-Q-F-G-R-D-I-A |
| 15 | B18 | 206 - 220 | K-H-T-P-I-N-L-V-R-D-L-P-Q-G-F |
| 16 | G15 | 791 - 805 | T-P-P-I-K-D-F-G-G-F-N-F-S-Q-I |
| 17 | H17 | 921 - 935 | K-L-I-A-N-Q-F-N-S-A-I-G-K-I-Q |
| 18 | J21 | 1181 - 1195 | K-E-I-D-R-L-N-E-V-A-K-N-L-N-E |
| 19 | H16 | 916 - 930 | L-Y-E-N-Q-K-L-I-A-N-Q-F-N-S-A |
| 20 | K13 | 1261 - 1275 | S-E-P-V-L-K-G-V-K-L-H-Y-T |

| #2 | Epitope ID | Position | Amino Acid Sequence |
| --- | --- | --- | --- |
| 1 | B11 | 171 - 185 | V-S-Q-P-F-L-M-D-L-E-G-K-Q-G-N |
| 2 | B12 | 176 - 190 | L-M-D-L-E-G-K-Q-G-N-F-K-N-L-R |
| 3 | A21 | 101 - 115 | I-R-G-W-I-F-G-T-T-L-D-S-K-T-Q |
| 4 | A10 | 46 - 60 | S-V-L-H-S-T-Q-D-L-F-L-P-F-F-S |
| 5 | G18 | 806 - 820 | L-P-D-P-S-K-P-S-K-R-S-F-I-E-D |
| 6 | A 1 | 1 - 15 | M-F-V-F-L-V-L-L-P-L-V-S-S-Q-C |
| 7 | A 5 | 21 - 35 | R-T-Q-L-P-P-A-Y-T-N-S-F-T-R-G |
| 8 | B 2 | 126 - 140 | V-V-I-K-V-C-E-F-Q-F-C-N-D-P-F |
| 9 | I 7 | 991 - 1005 | V-Q-I-D-R-L-I-T-G-R-L-Q-S-L-Q |
| 10 | F14 | 666 - 680 | I-G-A-G-I-C-A-S-Y-Q-T-Q-T-N-S |
| 11 | I 6 | 986 - 1000 | K-V-E-A-E-V-Q-I-D-R-L-I-T-G-R |
| 12 | A11 | 51 - 65 | T-Q-D-L-F-L-P-F-F-S-N-V-T-W-F |
| 13 | E20 | 576 - 590 | V-R-D-P-Q-T-L-E-I-L-D-I-T-P-C |
| 14 | E17 | 561 - 575 | P-F-Q-Q-F-G-R-D-I-A-D-T-T-D-A |
| 15 | D20 | 456 - 470 | F-R-K-S-N-L-K-P-F-E-R-D-I-S-T |
| 16 | B10 | 166 - 180 | C-T-F-E-Y-V-S-Q-P-F-L-M-D-L-E |
| 17 | E22 | 586 - 600 | D-I-T-P-C-S-F-G-G-V-S-V-I-T-P |
| 18 | H 5 | 861 - 875 | L-P-P-L-L-T-D-E-M-I-A-Q-Y-T-S |
| 19 | C 6 | 266 - 280 | Y-V-G-Y-L-Q-P-R-T-F-L-L-K-Y-N |
| 20 | A18 | 86 - 100 | F-N-D-G-V-Y-F-A-S-T-E-K-S-N-I |

| #3 | Epitope ID | Position | Amino Acid Sequence |
| --- | --- | --- | --- |
| 1 | E20 | 576 - 590 | V-R-D-P-Q-T-L-E-I-L-D-I-T-P-C |
| 2 | G20 | 816 - 830 | S-F-I-E-D-L-L-F-N-K-V-T-L-A-D |
| 3 | G19 | 811 - 825 | K-P-S-K-R-S-F-I-E-D-L-L-F-N-K |
| 4 | A 9 | 41 - 55 | K-V-F-R-S-S-V-L-H-S-T-Q-D-L-F |
| 5 | C14 | 306 - 320 | F-T-V-E-K-G-I-Y-Q-T-S-N-F-R-V |
| 6 | J14 | 1146 - 1160 | D-S-F-K-E-E-L-D-K-Y-F-K-N-H-T |
| 7 | K12 | 1256 - 1270 | F-D-E-D-D-S-E-P-V-L-K-G-V-K-L |
| 8 | E16 | 556 - 570 | N-K-K-F-L-P-F-Q-Q-F-G-R-D-I-A |
| 9 | E24 | 596 - 610 | S-V-I-T-P-G-T-N-T-S-N-Q-V-A-V |
| 10 | A16 | 76 - 90 | T-K-R-F-D-N-P-V-L-P-F-N-D-G-V |
| 11 | E19 | 571 - 585 | D-T-T-D-A-V-R-D-P-Q-T-L-E-I-L |
| 12 | A23 | 111 - 125 | D-S-K-T-Q-S-L-L-I-V-N-N-A-T-N |
| 13 | E15 | 551 - 565 | V-L-T-E-S-N-K-K-F-L-P-F-Q-Q-F |
| 14 | E23 | 591 - 605 | S-F-G-G-V-S-V-I-T-P-G-T-N-T-S |
| 15 | A15 | 71 - 85 | S-G-T-N-G-T-K-R-F-D-N-P-V-L-P |
| 16 | F 5 | 621 - 635 | P-V-A-I-H-A-D-Q-L-T-P-T-W-R-V |
| 17 | K11 | 1251 - 1265 | G-S-C-C-K-F-D-E-D-D-S-E-P-V-L |
| 18 | B21 | 221 - 235 | S-A-L-E-P-L-V-D-L-P-I-G-I-N-I |
| 19 | C 1 | 241 - 255 | L-L-A-L-H-R-S-Y-L-T-P-G-D-S-S |
| 20 | H11 | 891 - 905 | G-A-A-L-Q-I-P-F-A-M-Q-M-A-Y-R |

| #4 | Epitope ID | Position | Amino Acid Sequence |
| --- | --- | --- | --- |
| 1 | B21 | 221 - 235 | S-A-L-E-P-L-V-D-L-P-I-G-I-N-I |
| 2 | B 5 | 141 - 155 | L-G-V-Y-Y-H-K-N-N-K-S-W-M-E-S |
| 3 | B12 | 176 - 190 | L-M-D-L-E-G-K-Q-G-N-F-K-N-L-R |
| 4 | E15 | 551 - 565 | V-L-T-E-S-N-K-K-F-L-P-F-Q-Q-F |
| 5 | A 9 | 41 - 55 | K-V-F-R-S-S-V-L-H-S-T-Q-D-L-F |
| 6 | G11 | 771 - 785 | A-V-E-Q-D-K-N-T-Q-E-V-F-A-Q-V |
| 7 | I 3 | 971 - 985 | G-A-I-S-S-V-L-N-D-I-L-S-R-L-D |
| 8 | E14 | 546 - 560 | L-T-G-T-G-V-L-T-E-S-N-K-K-F-L |
| 9 | G10 | 766 - 780 | A-L-T-G-I-A-V-E-Q-D-K-N-T-Q-E |
| 10 | G19 | 811 - 825 | K-P-S-K-R-S-F-I-E-D-L-L-F-N-K |
| 11 | B 4 | 136 - 150 | C-N-D-P-F-L-G-V-Y-Y-H-K-N-N-K |
| 12 | G20 | 816 - 830 | S-F-I-E-D-L-L-F-N-K-V-T-L-A-D |
| 13 | H 1 | 841 - 855 | L-G-D-I-A-A-R-D-L-I-C-A-Q-K-F |
| 14 | J14 | 1146 - 1160 | D-S-F-K-E-E-L-D-K-Y-F-K-N-H-T |
| 15 | J22 | 1186 - 1200 | L-N-E-V-A-K-N-L-N-E-S-L-I-D-L |
| 16 | B13 | 181 - 195 | G-K-Q-G-N-F-K-N-L-R-E-F-V-F-K |
| 17 | G12 | 776 - 790 | K-N-T-Q-E-V-F-A-Q-V-K-Q-I-Y-K |
| 18 | H 2 | 846 - 860 | A-R-D-L-I-C-A-Q-K-F-N-G-L-T-V |
| 19 | H24 | 956 - 970 | A-Q-A-L-N-T-L-V-K-Q-L-S-S-N-F |
| 20 | F17 | 681 - 695 | P-R-R-A-R-S-V-A-S-Q-S-I-I-A-Y |

| #5 | Epitope ID | Position | Amino Acid Sequence |
| --- | --- | --- | --- |
| 1 | F22 | 706 - 720 | A-Y-S-N-N-S-I-A-I-P-T-N-F-T-I |
| 2 | E23 | 591 - 605 | S-F-G-G-V-S-V-I-T-P-G-T-N-T-S |
| 3 | B11 | 171 - 185 | V-S-Q-P-F-L-M-D-L-E-G-K-Q-G-N |
| 4 | F23 | 711 - 725 | S-I-A-I-P-T-N-F-T-I-S-V-T-T-E |
| 5 | E22 | 586 - 600 | D-I-T-P-C-S-F-G-G-V-S-V-I-T-P |
| 6 | F18 | 686 - 700 | S-V-A-S-Q-S-I-I-A-Y-T-M-S-L-G |
| 7 | F19 | 691 - 705 | S-I-I-A-Y-T-M-S-L-G-A-E-N-S-V |
| 8 | F21 | 701 - 715 | A-E-N-S-V-A-Y-S-N-N-S-I-A-I-P |
| 9 | B10 | 166 - 180 | C-T-F-E-Y-V-S-Q-P-F-L-M-D-L-E |
| 10 | J22 | 1186 - 1200 | L-N-E-V-A-K-N-L-N-E-S-L-I-D-L |
| 11 | H23 | 951 - 965 | V-V-N-Q-N-A-Q-A-L-N-T-L-V-K-Q |
| 12 | C16 | 316 - 330 | S-N-F-R-V-Q-P-T-E-S-I-V-R-F-P |
| 13 | F17 | 681 - 695 | P-R-R-A-R-S-V-A-S-Q-S-I-I-A-Y |
| 14 | J15 | 1151 - 1165 | E-L-D-K-Y-F-K-N-H-T-S-P-D-V-D |
| 15 | C10 | 286 - 300 | T-D-A-V-D-C-A-L-D-P-L-S-E-T-K |
| 16 | G 1 | 721 - 735 | S-V-T-T-E-I-L-P-V-S-M-T-K-T-S |
| 17 | E15 | 551 - 565 | V-L-T-E-S-N-K-K-F-L-P-F-Q-Q-F |
| 18 | F16 | 676 - 690 | T-Q-T-N-S-P-R-R-A-R-S-V-A-S-Q |
| 19 | K14 | 1266 - 1280 | E-P-V-L-K-G-V-K-L-H-Y-T |
| 20 | E21 | 581 - 595 | T-L-E-I-L-D-I-T-P-C-S-F-G-G-V |

| #6 | Epitope ID | Position | Amino Acid Sequence |
| --- | --- | --- | --- |
| 1 | G17 | 801 - 815 | N-F-S-Q-I-L-P-D-P-S-K-P-S-K-R |
| 2 | G15 | 791 - 805 | T-P-P-I-K-D-F-G-G-F-N-F-S-Q-I |
| 3 | G19 | 811 - 825 | K-P-S-K-R-S-F-I-E-D-L-L-F-N-K |
| 4 | G20 | 816 - 830 | S-F-I-E-D-L-L-F-N-K-V-T-L-A-D |
| 5 | E15 | 551 - 565 | V-L-T-E-S-N-K-K-F-L-P-F-Q-Q-F |
| 6 | K12 | 1256 - 1270 | F-D-E-D-D-S-E-P-V-L-K-G-V-K-L |
| 7 | G16 | 796 - 810 | D-F-G-G-F-N-F-S-Q-I-L-P-D-P-S |
| 8 | E19 | 571 - 585 | D-T-T-D-A-V-R-D-P-Q-T-L-E-I-L |
| 9 | E16 | 556 - 570 | N-K-K-F-L-P-F-Q-Q-F-G-R-D-I-A |
| 10 | G10 | 766 - 780 | A-L-T-G-I-A-V-E-Q-D-K-N-T-Q-E |
| 11 | J14 | 1146 - 1160 | D-S-F-K-E-E-L-D-K-Y-F-K-N-H-T |
| 12 | E20 | 576 - 590 | V-R-D-P-Q-T-L-E-I-L-D-I-T-P-C |
| 13 | J21 | 1181 - 1195 | K-E-I-D-R-L-N-E-V-A-K-N-L-N-E |
| 14 | K14 | 1266 - 1280 | E-P-V-L-K-G-V-K-L-H-Y-T |
| 15 | K13 | 1261 - 1275 | S-E-P-V-L-K-G-V-K-L-H-Y-T |
| 16 | D10 | 406 - 420 | E-V-R-Q-I-A-P-G-Q-T-G-K-I-A-D |
| 17 | J17 | 1161 - 1175 | S-P-D-V-D-L-G-D-I-S-G-I-N-A-S |
| 18 | J13 | 1141 - 1155 | L-Q-P-E-L-D-S-F-K-E-E-L-D-K-Y |
| 19 | G14 | 786 - 800 | K-Q-I-Y-K-T-P-P-I-K-D-F-G-G-F |
| 20 | C16 | 316 - 330 | S-N-F-R-V-Q-P-T-E-S-I-V-R-F-P |

| #7 | Epitope ID | Position | Amino Acid Sequence |
| --- | --- | --- | --- |
| 1 | G19 | 811 - 825 | K-P-S-K-R-S-F-I-E-D-L-L-F-N-K |
| 2 | A16 | 76 - 90 | T-K-R-F-D-N-P-V-L-P-F-N-D-G-V |
| 3 | F23 | 711 - 725 | S-I-A-I-P-T-N-F-T-I-S-V-T-T-E |
| 4 | G 1 | 721 - 735 | S-V-T-T-E-I-L-P-V-S-M-T-K-T-S |
| 5 | G18 | 806 - 820 | L-P-D-P-S-K-P-S-K-R-S-F-I-E-D |
| 6 | F24 | 716 - 730 | T-N-F-T-I-S-V-T-T-E-I-L-P-V-S |
| 7 | A17 | 81 - 95 | N-P-V-L-P-F-N-D-G-V-Y-F-A-S-T |
| 8 | G20 | 816 - 830 | S-F-I-E-D-L-L-F-N-K-V-T-L-A-D |
| 9 | G17 | 801 - 815 | N-F-S-Q-I-L-P-D-P-S-K-P-S-K-R |
| 10 | I 4 | 976 - 990 | V-L-N-D-I-L-S-R-L-D-K-V-E-A-E |
| 11 | E13 | 541 - 555 | F-N-F-N-G-L-T-G-T-G-V-L-T-E-S |
| 12 | A15 | 71 - 85 | S-G-T-N-G-T-K-R-F-D-N-P-V-L-P |
| 13 | B21 | 221 - 235 | S-A-L-E-P-L-V-D-L-P-I-G-I-N-I |
| 14 | I 2 | 966 - 980 | L-S-S-N-F-G-A-I-S-S-V-L-N-D-I |
| 15 | J22 | 1186 - 1200 | L-N-E-V-A-K-N-L-N-E-S-L-I-D-L |
| 16 | I15 | 1031 - 1045 | E-C-V-L-G-Q-S-K-R-V-D-F-C-G-K |
| 17 | I 1 | 961 - 975 | T-L-V-K-Q-L-S-S-N-F-G-A-I-S-S |
| 18 | C20 | 336 - 350 | C-P-F-G-E-V-F-N-A-T-R-F-A-S-V |
| 19 | E16 | 556 - 570 | N-K-K-F-L-P-F-Q-Q-F-G-R-D-I-A |
| 20 | B17 | 201 - 215 | F-K-I-Y-S-K-H-T-P-I-N-L-V-R-D |

## Discussion

Here, we report a screening and validation of predicted B cell epitopes of SARS-CoV-2 utilizing human serum from convalescing COVID-19 patients. We focused on S protein, especially on the RBD, because it has been reported that anti-RBD antibodies correlate well with an increase in spike-specific CD4^+^ T cell responses.

In the present study, patient sera were obtained from two different hospitals. Osaka University Hospital primarily admits severe patients requiring the ICU, and patient status might be in the subacute phase. Juso Osaka City Hospital, in contrast, usually admits mild or moderate patients, and patient status might be in the convalescent phase. Interestingly, the average antibody titer to S protein was higher in samples from Osaka University Hospital, which was consistent with the high titers in severe patients. Of importance, several patients possessed neutralizing activity with a high titer of IgG for S protein, which may suggest the functional importance of IgG for S protein as neutralizing antibodies. In addition, based on previous findings (Grifoni *et al*, 2020b), these results also suggest that the antibody titer to the RBD of S protein may predict an increase in spike-specific CD4^+^ T cell responses.

Antibodies targeting the RBD and S protein have enhanced potential for providing cross-protective immunity. The bioinformatics approach has been rapidly reported to identify potential B and T cell epitopes in S protein, which has provided data regarding antigen presentation, antibody-binding properties, and the predicted evolution of epitopes (Ahmed *et al*, 2020; Baruah & Bose, 2020; Bhattacharya *et al*, 2020; Grifoni *et al*, 2020a; Zheng & Song, 2020). A list of glycoprotein amino acid positions having a high probability of predicted B cell epitopes has been compiled. Based on the location of the relevant amino acid positions in the model structure, several epitope regions were predicted, i.e., 491-505 aa. and 558-562 aa. in the RBD, and the calculated surface of the amino acid residues of B cell epitopes are shown, i.e., 491-505 aa. and 558-562 aa. in the RBD and 1140-1146 aa. in other regions. We also performed predictions for linear B cell epitopes by BepiPred 2.0, the characterization of amino acids and the predicted structure. Seven regions (346-365 aa., 413-432 aa., 442-460 aa., 491-509 aa., and 518-537 aa. in the RBD and 671-690 aa. and 1146-1164 aa. in other regions) were prepared as B cell epitopes, which partially overlapped with previous reports (Ahmed *et al*, 2020; Baruah & Bose, 2020; Bhattacharya *et al*, 2020; Grifoni *et al*, 2020a; Zheng & Song, 2020). In this study, antibodies against the RBD were not evident, and only a few antibodies recognized these B cell epitopes in the RBD. These results indicate that the RBD region of S protein is not highly immunogenic, and the neutralizing antibodies against the RBD region may not be evident in individuals with COVID-19.

In this study, we evaluated linear B cell epitopes within nucleocapsid protein, membrane protein and envelope protein using a CelluSpots peptide array. Consistent with previous findings (Chow *et al*, 2006), most of the strongly binding B cell epitopes were located in nucleocapsid protein. These results suggest that nucleocapsid protein may be more immunogenic than S protein. However, a detailed analysis of antibody production by peptide array for S protein showed us possible candidate antigens in addition to the RBD. A few individuals (#2, #3, and #4) possessed neutralizing antibodies and showed several strongly binding epitopes in the NTD of S protein. Interestingly, Chie *et al* (2020) recently reported that a neutralizing human antibody binds to the NTD of S protein of SARS-CoV-2 but does not block the interaction between ACE2 and S protein. In their structural model, the monoclonal antibody interacts with the five loops for the NTD, especially between N3 (141-156 aa.) and N5 (246-260 aa.), and three glycosylation sites (Asn17, Asn61, and Asn149) were identified in this structure. Interestingly, as shown in Table 3, several epitopes in #2, #3, and #4 overlapped these regions in the NTD. Although our predicted epitope in the NTD (AH-528; 146-164 aa) did not react with the sera from the individuals in this study, we speculate that the NTD in S protein may be another candidate region for neutralizing antibodies.

As a study limitation, this study protocol has been approved to analyze only human serum samples without any clinical information. Because the onset of infection or severity of patients cannot be known, we cannot discuss the time course of antibodies with the clinical status of the patients. Although the magnitude of IgG production might be dependent on the duration of COVID-19, we can evaluate the dominant B cell epitope of each patient. There have been concerns regarding vaccine enhancement of disease by certain candidate COVID-19 vaccine approaches via antibody-dependent enhancement (ADE). This phenomenon is observed when non-neutralizing virus-specific IgG facilitates entry of virus particles into Fc-receptor-expressing cells, leading to inflammatory activation of macrophages and monocytes (Taylor *et al*, 2015). A study in SARS-CoV-1-infected *rhesus macaques* found that anti-S IgG contributes to severe acute lung injury (ALI) and massive accumulation of monocytes/macrophages in the lung (Liu *et al*, 2019). To avoid this phenomenon, a specific neutralizing antibody is required, which can also be achieved by a B cell epitope vaccine. Therefore, we speculate that the next generation of vaccines for COVID-19 should consider the B cell epitope in terms of safety and efficiency.

In summary, we conducted full B cell epitope mapping and validated the predicted B cell epitope of S protein, utilizing human sera from patients with COVID-19. Based on the analysis of neutralizing activity, anti-S antibodies might be correlated with the neutralizing action of the antibodies. The results may provide a novel target for the vaccine development against SARS-CoV-2.

## Materials and Methods

### Production of pseudotyped VSVs with S protein and transfection experiments

Pseudotyped vesicular stomatitis viruses (VSVs) and recombinant VSVs in which the G gene is replaced by a foreign reporter gene, such as luciferase, were generated. Either 293T or BHK cells were grown to 90% confluence on 35-mm tissue culture plates. The cells were infected with a recombinant vaccinia virus encoding the bacteriophage T7 RNA polymerase (vTF7-3) at a multiplicity of infection (MOI) of 5. After incubation at room temperature for 1 h, the cells were transfected with helper plasmids, pBS-N, pBS-P, pBS-L, and pBS-G, and template plasmids, pVSVΔG–Luci, using a cationic liposome reagent. After 4 h, the supernatants were replaced with 10% FBS DMEM, and the cells were incubated at 37°C for 48 h. The supernatants were then filtered through a 0.22-μm pore-size filter to remove vaccinia virus and were applied to 293T or BHK cells that had been transfected with pCAGVSVG 24 h previously. Recovery of the virus was assessed by examining the cells for the cytopathic effects that are typical of a VSV infection after 24 h. Stocks of *G-complemented viruses, i.e., VSVΔG virus or recombinant viruses transiently bearing VSV G protein on the virion surface, were grown from a single plaque on BHK cells transfected with pCAGVSVG and then stored at −80°C. The infectious titers of the recovered viruses were determined by a plaque assay. To generate pseudotype virus, 293T, BHK, or some other type of cells that exhibit a high competency of transfection were transfected with a plasmid expressing the envelope protein using a cationic liposome reagent. After 24 h of incubation at 37°C, cells were infected at an MOI of 0.5 with *G-VSVΔG–Luci. The virus was adsorbed for 2 h at 37°C and then extensively washed four times or more with serum-free DMEM. After 24 h of incubation at 37°C, the culture supernatants were collected, centrifuged to remove cell debris, and stored at −80°C. To generate pseudotype VSVs bearing the SARS-CoV-2 S protein, we transfected an expression plasmid encoding SARS-CoV-2 S protein. The pseudotype VSVs was prepared as described previously (Tani *et al*, 2007). Pseudotyped particles were harvested 24 h post-inoculation and clarified from cellular debris by centrifugation. For transfection, target cells were grown in 96-well plates until they reached 50-75% confluency before they were inoculated with the respective pseudotyped VSV.

### Serum samples

The serum samples of COVID-19 patients were obtained from Osaka University Hospital and Osaka City Juso Hospital. The protocol was approved by the ethical committee of Osaka University Hospital (No. 19546). Control serum was obtained from pooled human serum (#BJ11787, Tennessee Blood Services, TN, USA) mixed from five males and five females, which were collected in February 2019 and confirmed to be noninfected.

### Antibody test kits

SARS-CoV-2-specific IgG and IgM antibodies of serum samples collected from COVID-19 patients were detected using a rapid immunochromatographic test for detecting antibodies against SARS-CoV-2 (IgM; RF-NC001, IgG; RF-NC002, KURABO, Japan), a 2019-nCoV IgG/IgM detection kit (C6603C, Vazyme, China), a COVID-19 Human IgM/IgG Rapid Test (DC0301, Abnova, Taiwan), and a coronavirus (COVID-19) IgM/IgG rapid test kit (CG-CoV-IgM/IgG, RayBiotech, GA, USA), according to the manufacturers’ instructions.

### Synthetic SARS-CoV-2 peptides

Based on high antigenicity analysis of the three-dimensional predicted structure and B cell epitope information (BepiPred-2.0), eight different antigenic peptides were selected from the amino acid sequence of SARS-CoV-2 (Fig 3A and Appendix Table S2). The synthetic peptide was purified by reverse-phase HPLC (>98% purity) (Peptide Institute Inc., Osaka, Japan.). The synthetic SARS-CoV-2 peptide was reconstituted at 5 mg/ml in sterile PBS and stored below −20°C.

### Enzyme-linked immunosorbent assay

SARS-CoV-2 recombinant proteins, such as Recombinant 2019-nCoV Spike S1+S2 Protein (ECD; Beta Lifescience), Recombinant 2019-nCoV Spike Protein (RBD; Beta Lifescience), and other recombinant proteins (Appendix Table S1), were diluted in 50 mM carbonate buffer to a concentration of 1 μg/ml and coated at 50 ng/well on 96-well ELISA plates (MaxiSorp Nunc, Thermo Fisher Scientific K.K., Japan) overnight at 4°C. Synthetic SARS-CoV-2 peptides (Appendix Table S2; Peptide Institute Inc., Osaka, Japan) were diluted to a concentration of 10 μg/ml and coated at 500 ng/well. After blocking with 100 μl of sheep serum (16070-096, Gibco), the sera from COVID-19 patients were serially diluted from 10- to 31,250-fold in blocking buffer, added to each well, and incubated overnight at 4°C. After washing each well with 0.05% PBS Tween-20 (PBS-T), the cells were incubated with horseradish peroxidase (HRP)-conjugated antibodies specific for human IgG (1:10,000; AP004, Binding Site, U.K.) or IgM (1:5,000; AP012, Binding Site) with PBS containing 5% skim milk for 3 h at room temperature. For the IgG subclass determination assay, anti-human IgG subclass-specific HRP-conjugated antibodies (1:10,000; IgG1 (AP006), IgG2 (AP007), IgG3 (AP008), and IgG4 (AP009), Binding Site) were used. After washing the wells with PBS-T, color was developed with the peroxidase chromogenic substrate 3,3’,5,5’-tetramethylbenzidine (TMB; Sigma Aldrich, MO, USA), and the reactions were terminated with 0.9 N sulfuric acid. The absorbance was measured at 450 nm using an iMark microplate absorbance reader (Bio-Rad, CA, USA) and analyzed with MPM6 version 6.1 (Bio-Rad). The half-maximal antibody titer was analyzed with Prism GraphPad version 6.07 (GraphPad Software) and determined according to the highest value in the dilution range of each sample.

### Peptide array

CelluSpots peptide array was performed using CelluSpots™ covid19_hullB (98.301, Intavis) or CelluSpots™ covid19_hullS (98.302, Intavis) according to the manufacturers’ instructions: blocked by immersing the slides in PBS containing 5% skim milk overnight at 4°C on an orbital shaker, incubated with diluted serum samples (1:10) overnight at 4°C on an orbital shaker, washed with PBS-T buffer with 0.05% Tween-20, and incubated with diluted HRP-conjugated antibodies specific for human IgG (1:10,000; AP004, Binding Site) for 3 h at room temperature. A chemiluminescent signal visualized by Chemi-Lumi One L (07880-70, Nacalai Tesque) was detected with a ChemiDoc Touch imaging system (Bio-Rad) and analyzed with Image Lab software version 6.0.1 (Bio-Rad).

### Statistical analyses

All values are presented as the mean ± SEM. The statistical significance of differences between two groups was assessed by two-tailed unpaired t-test. A difference was considered statistically significant when p < 0.05. Statistical analysis was performed using Prism GraphPad version 6.07 (GraphPad Software).

## Acknowledgments

This study was supported by the Project Promoting Support for Drug Discovery grants (JP20nk0101602) from the Japan Agency for Medical Research and Development. We thank all members of the Department of Health Development and Medicine for supporting this project.

## Author contribution

S.Y. and H.N. wrote the manuscript. S.Y., S.S. K.T., H.A., Y.M. and H.N. designed the study with discussion. S.Y., C.O., and H.H performed the experiments and analyzed the data.

## The paper explained

### Problem

To fight against COVID-19, the rapid development of a vaccine is required. For this puroise, it is important to understand adaptive immunity to SARS-CoV-2 through the analysis of antibody production in coronavirus disease 2019 (COVID-19) patients. The spike glycoprotein (S) protein has been found to induce robust and protective humoral and cellular immunity, including the development of neutralizing antibodies and T cell-mediated immunity. We obtained serum from thirteen COVID-19 patients in Osaka University Hospital and Juso Osaka City Hospital who were diagnosed by RT-PCR for SARS-CoV-2 infection. Here, we addressed the humoral immune response by measuring antibody production against S protein and the neutralizing ability in convalescent patients from two different hospitals.

### Results

We analyzed the neutralizing activities against SARS-CoV-2 assessed by a pseudotype virus-neutralizing assay. Most individuals revealed neutralizing activity; however, they were not observed in a few patients. The antibody production against the spike glycoprotein (S protein) or receptor-binding domain (RBD) of SARS-CoV-2 was elevated, with large individual differences, as assessed by ELISA. The titer for anti-S IgG in patients from Osaka University Hospital was relatively higher than that of anti-RBD antibodies in patients from Juso Osaka City Hospital. However, the titer for anti-RBD antibodies was higher in patients from Juso Osaka City Hospital. As most of the patients in Osaka University Hospital were cared for in the intensive care unit, the disease phase or severity may affect their titers. We also predicted the linear B cell epitopes assessed by the characterization of the predicted amino acid structure. Two regions (671-690 aa. and 1146-1164 aa.), which were located in S1 and S2 but not in the RBD, were highly reactive with the sera from patients.

### Impact

The analysis of antibody production and B cell epitopes of the S protein from patient serum provided important information for the protective effect against SAR-CoV-2. Although the magnitude of IgG production might be dependent on the duration of COVID-19, we can evaluate the dominant B cell epitope of each patient. We further addressed the B cell epitope within the S protein by utilizing a B cell epitope array. Although there were large individual differences, a hot spot in the N-terminal domain of the S protein but not the RBD was observed in individuals with neutralizing activity. These results provided a novel target for the vaccine development against SARS-CoV-2.

