## Supplement for "SARS-CoV-2-induced humoral immunity through B cell epitope analysis and neutralizing activity in COVID-19 infected individuals in Japan"

### APPENDIX

Table of Contents

Page 2: Appendix Table S1. Candidate recombinant proteins for ELISA
Page 3: Appendix Table S2. Amino acid sequence of each synthetic SARS-CoV-2 peptide
Page 4,5: Appendix Table S3. Top 20 strongest binding peptide regions of serum samples from each individual (OU #1-7)
Page 6: Appendix Figure S1. Anti-SARS-CoV-2 IgG2 and IgG4 responses of COVID-19 patients.

#### SARS-CoV-2-induced humoral immunity through B cell epitope analysis and neutralizing activity in COVID-19 infected individuals in Japan

Shota Yoshida,[1, 2] Chikako Ono [3], Hiroki Hayashi [1],
Satoshi Shiraishi [4], Kazunori Tomono[5],
Hisashi Arase [6,7], Yoshiharu Matsuura[3], Hironori Nakagami [1*]

[1]Department of Health Development and Medicine, Osaka University Graduate School of Medicine.
[2]Department of Geriatric and General Medicine, Osaka University Graduate School of Medicine.
[3]Department of Molecular Virology, Research Institute for Microbial Diseases, Osaka University
[4]Juso Osaka City.Hospital
[5]Division of Infection Control and Prevention, Osaka University Hospital
[6]Department of Immunochemistry, Researsch Institute for Microbial Diseases, Osaka University, Suita, Osaka 565-0871, Japan
[7]Laboratory of Immunochemistry, WPI Immunology Frontier Research Centre, Osaka University, Suita, Osaka 565-0871

### Appendix Table S1.

#### Candidate recombinant proteins for ELISA

| Product | Manufacturer | Catalog Code | Expression System |
|---|---|---|---|
| SARS-CoV-2 (2019-nCoV) Spike S1-His Recombinant Protein | Sino Biological | 40591-V08H | HEK293 Cells |
| SARS-CoV-2 (2019-nCoV) Spike RBD-Fc Recombinant Protein | Sino Biological | 40592-V02H | HEK293 Cells |
| Recombinant 2019-nCoV/COVID-19 Spike S1 Protein (His Tag) | Beta Lifescience | BLPSN-0982P | HEK293 Cells |
| Recombinant 2019-nCoV Spike S1+S2 Protein (ECD, His tag) | Beta Lifescience | BLPSN-0986P | Baculovirus-Insect Cells |
| Recombinant 2019-nCoV Spike Protein S2 (ECD, His tag) | Beta Lifescience | BLPSN-0987P | Baculovirus-Insect Cells |
| Recombinant 2019-nCoV Spike Protein (RBD, His Tag) | Beta Lifescience | BLPSN-0988P | Baculovirus-Insect Cells |
| Recombinant SARS-CoV-2 Spike Protein, S1 Subunit | Ray Biotech | 230-01101 | Escherichia coli |
| Recombinant SARS-CoV-2 Spike Protein, S2 Subunit | Ray Biotech | 230-01103 | Escherichia coli |

### Appendix Table S2.

#### Amino acid sequence of each synthetic SARS-CoV-2 peptide

| Peptide ID | Position | Amino Acid Sequence |
|---|---|---|
| AH-526 | 413 - 432 | G-Q-T-G-K-I-A-D-Y-N-Y-K-L-P-D-D-F-T-G-C |
| AH-527 | 442 - 460 | D-S-K-V-G-G-N-Y-N-Y-L-Y-R-L-F-R-K-S-N |
| AH-528 | 146 - 164 | H-K-N-N-K-S-W-M-E-S-E-F-R-V-Y-S-S-A-N |
| AH-529 | 671 - 690 | C-A-S-Y-Q-T-Q-T-N-S-P-R-R-A-R-S-V-A-S-Q |
| AH-530 | 1146 - 1164 | D-S-F-K-E-E-L-D-K-Y-F-K-N-H-T-S-P-D-V |
| AH-531 | 346 - 365 | R-F-A-S-V-Y-A-W-N-R-K-R-I-S-N-C-V-A-D-Y |
| AH-532 | 491 - 509 | P-L-Q-S-Y-G-F-Q-P-T-N-G-V-G-Y-Q-P-Y-R |
| AH-533 | 518 - 537 | L-H-A-P-A-T-V-C-G-P-K-K-S-T-N-L-V-K-N-K |

### Appendix Table S3.

#### Top 20 strongest binding peptide regions of serum samples from each individual (OU #1-7)

| #1 | Epitope ID | Type | Position | Amino Acid Sequence |
|---|---|---|---|---|
| 1 | A4 | N | 10 - 24 | R-N-A-P-R-I-T-F-G-G-P-S-D-S-T |
| 2 | A5 | N | 13 - 27 | P-R-I-T-F-G-G-P-S-D-S-T-G-S-N |
| 3 | A3 | N | 7 - 21 | Q-N-Q-R-N-A-P-R-I-T-F-G-G-P-S |
| 4 | A2 | N | 4 - 18 | N-G-P-Q-N-Q-R-N-A-P-R-I-T-F-G |
| 5 | F13 | N | 397 - 411 | A-A-D-L-D-D-F-S-K-Q-L-Q-Q-S-M |
| 6 | C24 | N | 214 - 228 | G-G-D-A-A-L-A-L-L-L-L-D-R-L-N |
| 7 | J23 | M | 211 - 225 | S-S-S-S-D-N-I-A-L-L-V-Q |
| 8 | A12 | N | 34 - 48 | G-A-R-S-K-Q-R-R-P-Q-G-L-P-N-N |
| 9 | A13 | N | 37 - 51 | S-K-Q-R-R-P-Q-G-L-P-N-N-T-A-S |
| 10 | H3 | M | 7 - 21 | T-I-T-V-E-E-L-K-K-L-L-E-Q-W-N |
| 11 | B18 | N | 124 - 138 | G-A-N-K-D-G-I-I-W-V-A-T-E-G-A |
| 12 | D2 | N | 220 - 234 | A-L-L-L-L-D-R-L-N-Q-L-E-S-K-M |
| 13 | C23 | N | 211 - 225 | A-G-N-G-G-D-A-A-L-A-L-L-L-L-D |
| 14 | H2 | M | 4 - 18 | S-N-G-T-I-T-V-E-E-L-K-K-L-L-E |
| 15 | J22 | M | 208 - 222 | T-D-H-S-S-S-S-D-N-I-A-L-L-V-Q |
| 16 | F8 | N | 382 - 396 | L-P-Q-R-Q-K-K-Q-Q-T-V-T-L-L-P |
| 17 | D1 | N | 217 - 231 | A-A-L-A-L-L-L-L-D-R-L-N-Q-L-E |
| 18 | B19 | N | 127 - 141 | K-D-G-I-I-W-V-A-T-E-G-A-L-N-T |
| 19 | F14 | N | 400 - 414 | L-D-D-F-S-K-Q-L-Q-Q-S-M-S-S-A |
| 20 | B17 | N | 121 - 135 | L-P-Y-G-A-N-K-D-G-I-I-W-V-A-T |

| #2 | Epitope ID | Type | Position | Amino Acid Sequence |
|---|---|---|---|---|
| 1 | C1 | N | 145 - 159 | H-I-G-T-R-N-P-A-N-N-A-A-I-V-L |
| 2 | D2 | N | 220 - 234 | A-L-L-L-L-D-R-L-N-Q-L-E-S-K-M |
| 3 | C2 | N | 148 - 162 | T-R-N-P-A-N-N-A-A-I-V-L-Q-L-P |
| 4 | D1 | N | 217 - 231 | A-A-L-A-L-L-L-L-D-R-L-N-Q-L-E |
| 5 | A4 | N | 10 - 24 | R-N-A-P-R-I-T-F-G-G-P-S-D-S-T |
| 6 | A6 | N | 16 - 30 | T-F-G-G-P-S-D-S-T-G-S-N-Q-N-G |
| 7 | A5 | N | 13 - 27 | P-R-I-T-F-G-G-P-S-D-S-T-G-S-N |
| 8 | J14 | M | 184 - 198 | S-Q-R-V-A-G-D-S-G-F-A-A-Y-S-R |
| 9 | A7 | N | 19 - 33 | G-P-S-D-S-T-G-S-N-Q-N-G-E-R-S |
| 10 | C5 | N | 157 - 171 | I-V-L-Q-L-P-Q-G-T-T-L-P-K-G-F |
| 11 | D5 | N | 229 - 243 | Q-L-E-S-K-M-S-G-K-G-Q-Q-Q-Q-G |
| 12 | J11 | M | 175 - 189 | T-L-S-Y-Y-K-L-G-A-S-Q-R-V-A-G |
| 13 | A3 | N | 7 - 21 | Q-N-Q-R-N-A-P-R-I-T-F-G-G-P-S |
| 14 | A2 | N | 4 - 18 | N-G-P-Q-N-Q-R-N-A-P-R-I-T-F-G |
| 15 | J12 | M | 178 - 192 | Y-Y-K-L-G-A-S-Q-R-V-A-G-D-S-G |
| 16 | D4 | N | 226 - 240 | R-L-N-Q-L-E-S-K-M-S-G-K-G-Q-Q |
| 17 | D3 | N | 223 - 237 | L-L-D-R-L-N-Q-L-E-S-K-M-S-G-K |
| 18 | H3 | M | 7 - 21 | T-I-T-V-E-E-L-K-K-L-L-E-Q-W-N |
| 19 | J15 | M | 187 - 201 | V-A-G-D-S-G-F-A-A-Y-S-R-Y-R-I |
| 20 | F1 | N | 361 - 375 | K-T-F-P-P-T-E-P-K-K-D-K-K-K-K |

| #3 | Epitope ID | Type | Position | Amino Acid Sequence |
|---|---|---|---|---|
| 1 | C24 | N | 214 - 228 | G-G-D-A-A-L-A-L-L-L-L-D-R-L-N |
| 2 | C23 | N | 211 - 225 | A-G-N-G-G-D-A-A-L-A-L-L-L-L-D |
| 3 | D1 | N | 217 - 231 | A-A-L-A-L-L-L-L-D-R-L-N-Q-L-E |
| 4 | A22 | N | 64 - 78 | L-K-F-P-R-G-Q-G-V-P-I-N-T-N-S |
| 5 | A3 | N | 7 - 21 | Q-N-Q-R-N-A-P-R-I-T-F-G-G-P-S |
| 6 | D2 | N | 220 - 234 | A-L-L-L-L-D-R-L-N-Q-L-E-S-K-M |
| 7 | A4 | N | 10 - 24 | R-N-A-P-R-I-T-F-G-G-P-S-D-S-T |
| 8 | A2 | N | 4 - 18 | N-G-P-Q-N-Q-R-N-A-P-R-I-T-F-G |
| 9 | A23 | N | 67 - 81 | P-R-G-Q-G-V-P-I-N-T-N-S-S-P-D |
| 10 | C16 | N | 190 - 204 | S-R-N-S-S-R-N-S-T-P-G-S-S-R-G |
| 11 | A24 | N | 70 - 84 | Q-G-V-P-I-N-T-N-S-S-P-D-D-Q-I |
| 12 | B1 | N | 73 - 87 | P-I-N-T-N-S-S-P-D-D-Q-I-G-Y-Y |
| 13 | A14 | N | 40 - 54 | R-R-P-Q-G-L-P-N-N-T-A-S-W-F-T |
| 14 | C15 | N | 187 - 201 | S-S-R-S-R-N-S-S-R-N-S-T-P-G-S |
| 15 | F2 | N | 364 - 378 | P-P-T-E-P-K-K-D-K-K-K-K-A-D-E |
| 16 | C18 | N | 196 - 210 | N-S-T-P-G-S-S-R-G-T-S-P-A-R-M |
| 17 | F1 | N | 361 - 375 | K-T-F-P-P-T-E-P-K-K-D-K-K-K-K |
| 18 | A21 | N | 61 - 75 | K-E-D-L-K-F-P-R-G-Q-G-V-P-I-N |
| 19 | D3 | N | 223 - 237 | L-L-D-R-L-N-Q-L-E-S-K-M-S-G-K |
| 20 | F10 | N | 388 - 402 | K-Q-Q-T-V-T-L-L-P-A-A-D-L-D-D |

| #4 | Epitope ID | Type | Position | Amino Acid Sequence |
|---|---|---|---|---|
| 1 | A11 | N | 31 - 45 | E-R-S-G-A-R-S-K-Q-R-R-P-Q-G-L |
| 2 | A12 | N | 34 - 48 | G-A-R-S-K-Q-R-R-P-Q-G-L-P-N-N |
| 3 | A21 | N | 61 - 75 | K-E-D-L-K-F-P-R-G-Q-G-V-P-I-N |
| 4 | C2 | N | 148 - 162 | T-R-N-P-A-N-N-A-A-I-V-L-Q-L-P |
| 5 | A22 | N | 64 - 78 | L-K-F-P-R-G-Q-G-V-P-I-N-T-N-S |
| 6 | B24 | N | 142 - 156 | P-K-D-H-I-G-T-R-N-P-A-N-N-A-A |
| 7 | A20 | N | 58 - 72 | Q-H-G-K-E-D-L-K-F-P-R-G-Q-G-V |
| 8 | C1 | N | 145 - 159 | H-I-G-T-R-N-P-A-N-N-A-A-I-V-L |
| 9 | C7 | N | 163 - 177 | Q-G-T-T-L-P-K-G-F-Y-A-E-G-S-R |
| 10 | A13 | N | 37 - 51 | S-K-Q-R-R-P-Q-G-L-P-N-N-T-A-S |
| 11 | D2 | N | 220 - 234 | A-L-L-L-L-D-R-L-N-Q-L-E-S-K-M |
| 12 | F5 | N | 373 - 387 | K-K-K-A-D-E-T-Q-A-L-P-Q-R-Q-K |
| 13 | F6 | N | 376 - 390 | A-D-E-T-Q-A-L-P-Q-R-Q-K-K-Q-Q |
| 14 | C4 | N | 154 - 168 | N-A-A-I-V-L-Q-L-P-Q-G-T-T-L-P |
| 15 | C5 | N | 157 - 171 | I-V-L-Q-L-P-Q-G-T-T-L-P-K-G-F |
| 16 | F10 | N | 388 - 402 | K-Q-Q-T-V-T-L-L-P-A-A-D-L-D-D |
| 17 | C6 | N | 160 - 174 | Q-L-P-Q-G-T-T-L-P-K-G-F-Y-A-E |
| 18 | F11 | N | 391 - 405 | T-V-T-L-L-P-A-A-D-L-D-D-F-S-K |
| 19 | F13 | N | 397 - 411 | A-A-D-L-D-D-F-S-K-Q-L-Q-Q-S-M |
| 20 | D10 | N | 244 - 258 | Q-T-V-T-K-K-S-A-A-E-A-S-K-K-P |

N, Nucleocapsid; M, Membrane; E, Envelope.

### Appendix Table S3.

Top 20 strongest binding peptide regions of serum samples from each individual (OU #1-7)

| #5 | Epitope ID | Type | Position | Amino Acid Sequence |
|---|---|---|---|---|
| 1 | D2 | N | 220 - 234 | A-L-L-L-L-D-R-L-N-Q-L-E-S-K-M |
| 2 | C19 | N | 199 - 213 | P-G-S-S-R-G-T-S-P-A-R-M-A-G-N |
| 3 | C18 | N | 196 - 210 | N-S-T-P-G-S-S-R-G-T-S-P-A-R-M |
| 4 | D22 | N | 280 - 294 | E-Q-T-Q-G-N-F-G-D-Q-E-L-I-R-Q |
| 5 | C17 | N | 193 - 207 | S-S-R-N-S-T-P-G-S-S-R-G-T-S-P |
| 6 | B16 | N | 118 - 132 | E-A-G-L-P-Y-G-A-N-K-D-G-I-I-W |
| 7 | D18 | N | 268 - 282 | Y-N-V-T-Q-A-F-G-R-R-G-P-E-Q-T |
| 8 | B22 | N | 136 - 150 | E-G-A-L-N-T-P-K-D-H-I-G-T-R-N |
| 9 | D3 | N | 223 - 237 | L-L-D-R-L-N-Q-L-E-S-K-M-S-G-K |
| 10 | C23 | N | 211 - 225 | A-G-N-G-G-D-A-A-L-A-L-L-L-L-D |
| 11 | C15 | N | 187 - 201 | S-S-R-S-R-N-S-S-R-N-S-T-P-G-S |
| 12 | C24 | N | 214 - 228 | G-G-D-A-A-L-A-L-L-L-L-D-R-L-N |
| 13 | B1 | N | 73 - 87 | P-I-N-T-N-S-S-P-D-D-Q-I-G-Y-Y |
| 14 | J1 | M | 145 - 159 | L-R-G-H-L-R-I-A-G-H-H-L-G-R-C |
| 15 | L3 | E | 7 - 21 | E-E-T-G-T-L-I-V-N-S-V-L-L-F-L |
| 16 | I23 | M | 139 - 153 | V-I-G-A-V-I-L-R-G-H-L-R-I-A-G |
| 17 | H1 | M | 1 - 15 | M-A-D-S-N-G-T-I-T-V-E-E-L-K-K |
| 18 | E1 | N | 289 - 303 | Q-E-L-I-R-Q-G-T-D-Y-K-H-W-P-Q |
| 19 | J2 | M | 148 - 162 | H-L-R-I-A-G-H-H-L-G-R-C-D-I-K |
| 20 | D19 | N | 271 - 285 | T-Q-A-F-G-R-R-G-P-E-Q-T-Q-G-N |

| #6 | Epitope ID | Type | Position | Amino Acid Sequence |
|---|---|---|---|---|
| 1 | F16 | N | 406 - 420 | Q-L-Q-Q-S-M-S-S-A-D-S-T-Q-A |
| 2 | J22 | M | 208 - 222 | T-D-H-S-S-S-S-D-N-I-A-L-L-V-Q |
| 3 | F15 | N | 403 - 417 | F-S-K-Q-L-Q-Q-S-M-S-S-A-D-S-T |
| 4 | F2 | N | 364 - 378 | P-P-T-E-P-K-K-D-K-K-K-K-A-D-E |
| 5 | F17 | N | 409 - 423 | Q-Q-S-M-S-S-A-D-S-T-Q-A |
| 6 | J21 | M | 205 - 219 | K-L-N-T-D-H-S-S-S-S-D-N-I-A-L |
| 7 | A13 | N | 37 - 51 | S-K-Q-R-R-P-Q-G-L-P-N-N-T-A-S |
| 8 | A12 | N | 34 - 48 | G-A-R-S-K-Q-R-R-P-Q-G-L-P-N-N |
| 9 | A5 | N | 13 - 27 | P-R-I-T-F-G-G-P-S-D-S-T-G-S-N |
| 10 | D2 | N | 220 - 234 | A-L-L-L-L-D-R-L-N-Q-L-E-S-K-M |
| 11 | C21 | N | 205 - 219 | T-S-P-A-R-M-A-G-N-G-G-D-A-A-L |
| 12 | A14 | N | 40 - 54 | R-R-P-Q-G-L-P-N-N-T-A-S-W-F-T |
| 13 | E24 | N | 358 - 372 | D-A-Y-K-T-F-P-P-T-E-P-K-K-D-K |
| 14 | J23 | M | 211 - 225 | S-S-S-S-D-N-I-A-L-L-V-Q |
| 15 | B1 | N | 73 - 87 | P-I-N-T-N-S-S-P-D-D-Q-I-G-Y-Y |
| 16 | C15 | N | 187 - 201 | S-S-R-S-R-N-S-S-R-N-S-T-P-G-S |
| 17 | C14 | N | 184 - 198 | S-R-S-S-S-R-S-R-N-S-S-R-N-S-T |
| 18 | C24 | N | 214 - 228 | G-G-D-A-A-L-A-L-L-L-L-D-R-L-N |
| 19 | A11 | N | 31 - 45 | E-R-S-G-A-R-S-K-Q-R-R-P-Q-G-L |
| 20 | F8 | N | 382 - 396 | L-P-Q-R-Q-K-K-Q-Q-T-V-T-L-L-P |

| #7 | Epitope ID | Type | Position | Amino Acid Sequence |
|---|---|---|---|---|
| 1 | A22 | N | 64 - 78 | L-K-F-P-R-G-Q-G-V-P-I-N-T-N-S |
| 2 | A23 | N | 67 - 81 | P-R-G-Q-G-V-P-I-N-T-N-S-S-P-D |
| 3 | C24 | N | 214 - 228 | G-G-D-A-A-L-A-L-L-L-L-D-R-L-N |
| 4 | B24 | N | 142 - 156 | P-K-D-H-I-G-T-R-N-P-A-N-N-A-A |
| 5 | F9 | N | 385 - 399 | R-Q-K-K-Q-Q-T-V-T-L-L-P-A-A-D |
| 6 | A13 | N | 37 - 51 | S-K-Q-R-R-P-Q-G-L-P-N-N-T-A-S |
| 7 | F10 | N | 388 - 402 | K-Q-Q-T-V-T-L-L-P-A-A-D-L-D-D |
| 8 | B23 | N | 139 - 153 | L-N-T-P-K-D-H-I-G-T-R-N-P-A-N |
| 9 | F13 | N | 397 - 411 | A-A-D-L-D-D-F-S-K-Q-L-Q-Q-S-M |
| 10 | A20 | N | 58 - 72 | Q-H-G-K-E-D-L-K-F-P-R-G-Q-G-V |
| 11 | A21 | N | 61 - 75 | K-E-D-L-K-F-P-R-G-Q-G-V-P-I-N |
| 12 | A4 | N | 10 - 24 | R-N-A-P-R-I-T-F-G-G-P-S-D-S-T |
| 13 | D10 | N | 244 - 258 | Q-T-V-T-K-K-S-A-A-E-A-S-K-K-P |
| 14 | F8 | N | 382 - 396 | L-P-Q-R-Q-K-K-Q-Q-T-V-T-L-L-P |
| 15 | D12 | N | 250 - 264 | S-A-A-E-A-S-K-K-P-R-Q-K-R-T-A |
| 16 | F11 | N | 391 - 405 | T-V-T-L-L-P-A-A-D-L-D-D-F-S-K |
| 17 | D1 | N | 217 - 231 | A-A-L-A-L-L-L-L-D-R-L-N-Q-L-E |
| 18 | A3 | N | 7 - 21 | Q-N-Q-R-N-A-P-R-I-T-F-G-G-P-S |
| 19 | H2 | M | 4 - 18 | S-N-G-T-I-T-V-E-E-L-K-K-L-L-E |
| 20 | D3 | N | 223 - 237 | L-L-D-R-L-N-Q-L-E-S-K-M-S-G-K |

N, Nucleocapsid; M, Membrane; E, Envelope.

Appendix Figure S1.

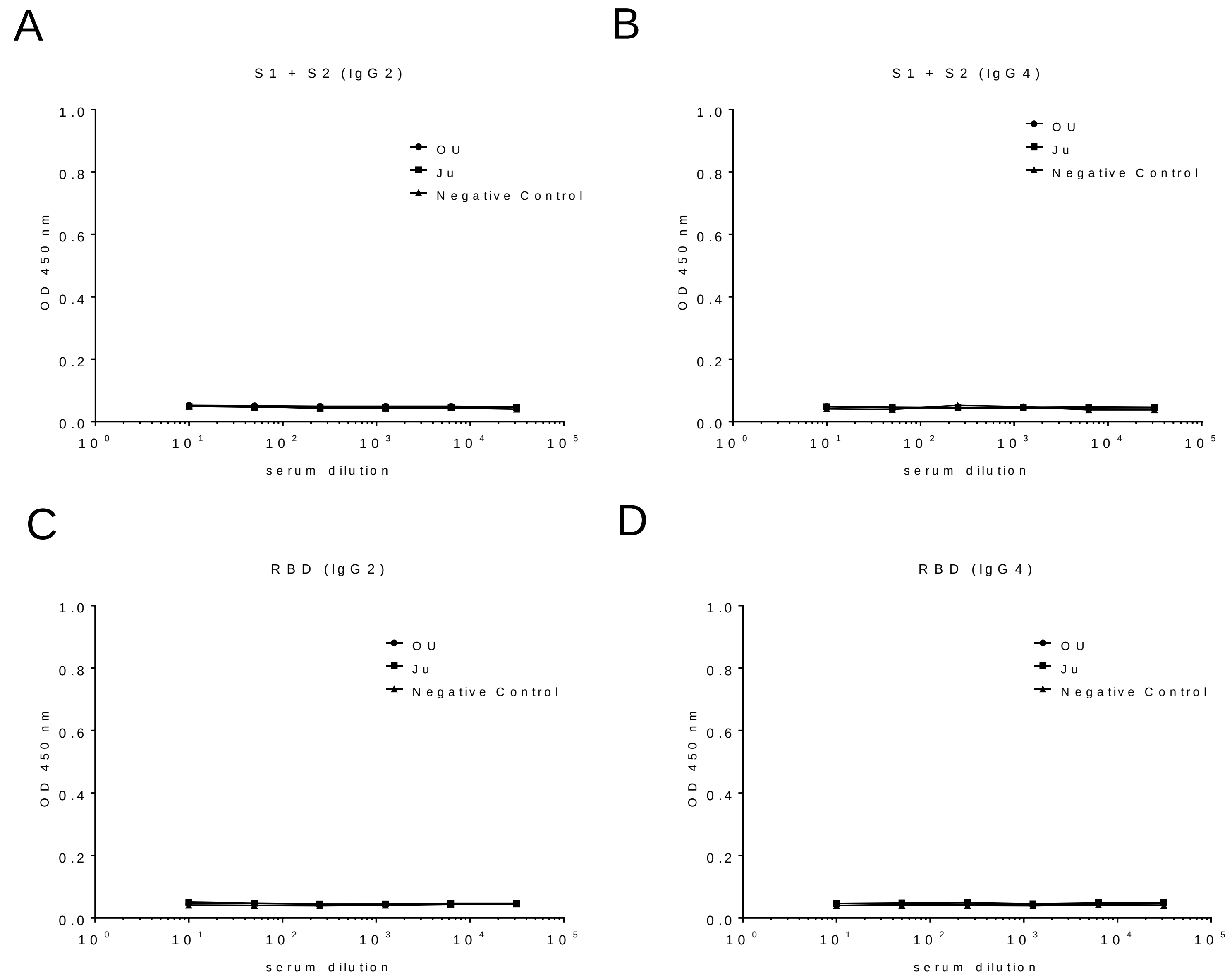


**Anti-SARS-CoV-2 IgG2 and IgG4 responses of COVID-19 patients.**

**(A and B)** The serum titer against recombinant SARS-CoV-2 spike S1+S2 protein. **(A)** IgG2, **(B)** IgG4, expressed as the OD at 450 nm. **(C and D)** The serum titer against recombinant SARS-CoV-2 spike RBD protein. **(C)** IgG2, **(D)** IgG4, expressed as the OD at 450 nm. OU, serum samples collected from patients in the ICU of Osaka University Hospital; Ju, serum samples collected from patients in Osaka City Juso Hospital. All the data are expressed as the mean ± SEM.
